# *In-vitro* evaluation of *Pseudomonas fluorescens* antibacterial activity against *Listeria* spp. isolated from New Zealand horticultural environments

**DOI:** 10.1101/2020.07.03.183277

**Authors:** Vathsala Mohan, Graham Fletcher, Françoise Leroi

## Abstract

Beneficial bacteria with antibacterial properties are an attractive alternative to chemical-based antibacterial or bactericidal agents. The aim of our study was to source such bacteria from horticultural produce and environments and to explore the mechanisms of their antimicrobial properties. Four strains of *Pseudomonas fluorescens* were isolated that possessed antibacterial activity against the pathogen *Listeria monocytogenes.* These strains (PFR46H06, PFR46H07, PFR46H08 and PFR46H09) were tested against *L. monocytogenes* (n=31), *L. seeligeri* (n=1) and *L. innocua* (n=1) isolated from seafood and horticultural sources, and two *L. monocytogenes* from clinical cases (Scott A and ATCC 49594, n=2). All *Listeria* strains were inhibited by the three strains PFR46H07, PFR46H08 and PFR46H09 with average zones of inhibition of 14.8, 15.1 and 18.2 mm, respectively, with PFR46H09 having significantly more inhibition than the other two (p<0.05). PFR46H06 showed minimal inhibition towards the *L. seeligeri* and *L. innocua* isolates and no inhibition towards most of the *L. monocytogenes* isolates. In order to investigate the functional protein differences between these strains, all four strains were subjected to whole cell lysate proteomics; data are available via ProteomeXchange with identifier PXD019965. We found significant differences in the peptide profiles and protein summaries between these isolates. The total number of proteins identified was 1781 in PFR46H06, 2030 in PFR46H07, 2228 in PFR46H08 and 1994 in PFR46H09. We paid special attention to secretion system proteins as these play a main role in the defense mechanisms of bacteria, particularly those of the type VI secretion system (T6SS) and found that they varied significantly (p<0.001) among the four isolates. PFR46H06 and PFR46H07 possessed the fewest secretion system structural proteins (12 and 11 respectively) while PFR46H08 and PFR46H09 each had 18. PFR46H09, which showed the greatest antimicrobial effect, had nine T6SS proteins compared to just four in the other three strains.

## Introduction

There is a strong demand for sustainable technologies to improve the quality and safety of fresh produce and food production. As part of established appropriate horticultural practices for the production and sale of safe food products, growers rely on physical washing and chemical sanitizers. Use of chemical-based sanitizers can pose health risks and cause undesirable environmental effects. This has resulted in increasing concern over current control measures and chemicals such as chlorine are being banned in some countries (Van Haute, Sampers et al. 2013). Alternative control measures for pathogens are therefore being sought, including other synthetic chemicals such as organophosphates, carbamates and pyrethroids (Aktar, Sengupta et al. 2009) but unfortunately controversies have been raised for the use of these chemicals and the other mode of pathogen control is to implement biocontrol measures. Biological control agents (BCAs) are living organisms that are used to control unwanted organisms. They have been used in different fields of biology, particularly in entomology and plant pathology against pests and microbial pathogens to suppress their populations (Weller 2007).

*Listeria monocytogenes* is a Gram positive, non-sporulating bacillus that is ubiquitous in nature and has been isolated from a wide range of sources including horticultural produce, processed foods, dairy products, silage and soils (Albano, Oliveira et al. 2007, Lakicevic, Nastasijevic et al. 2015). Being a facultative intracellular pathogen, *L. monocytogenes* can cause invasive diseases such as meningoencephalitis, sepsis, and gastroenteritis in immuno-compromised humans, as well as miscarriage in pregnant women. It causes disease in several farm animals including cows, sheep, pigs, and goats (Mohammed, Atwill et al. 2010, Buchrieser, Rusniok et al. 2011). Human disease occurs as a result of direct contact with infected animals or due to ingestion of contaminated food products (Dimitrijevic, Anderson et al. 2006, Oevermann, Zurbriggen et al. 2010, Schuppler and Loessner 2010).

Bio-preservation refers to the use of non-pathogenic microorganisms and/or their metabolites to extend the shelf-life of food and to improve the microbiological quality (García, Rodríguez et al. 2010, Gaggia, Di Gioia et al. 2011).In particular, lactic acid bacteria (LAB) are being used to aid food preservation. LABs have a long history of use in fermented foods, making them attractive choices to be used for bio-preservation, particularly for protection against *L. monocytogenes* (Leroi 2010).

*Pseudomonas fluorescens* comprises a group of saprophytes that commonly colonize soil, water and plant surface environments. *P. fluorescens* is a Gram-negative, rod-shaped bacterium that secretes a soluble greenish fluorescent pigment called fluorescein, particularly under conditions of low iron availability (Meyer and Abdallah 1978). Most *P. fluorescens* strains are obligate aerobes, although some strains take up NO_3_ for respiration in place of O_2_. The species is motile with multiple polar flagella. *P. fluorescens* has been demonstrated to grow well in mineral salts media with carbon sources (Palleroni 1984). *P. fluorescens* is a plant growth promoting rhizobacterium (PGPR) and has been identified as a potential BCA for bacterial diseases in plants (Sunita, Saleena et al. 2003, Weller 2007). Its saprophytic nature and natural soil adaptation abilities permit robust survival in soil. Certain strains have proven to be potent (BCAs that suppress plant diseases by protecting the seeds and roots from fungal infection (Hoffland, Hakulinen et al. 1996). *P. fluorescens* produces secondary metabolites including antibiotics, siderophores and hydrogen cyanide (O’Sullivan and O’Gara 1992). A review by Haas and Defago (2005), identified rapid colonization through competitive exclusion of pathogens as the main mechanism of action by which *P. fluorescens* inhibits pathogens in the rhizosphere. Although *P. fluorescens* is not generally considered a bacterial pathogen in humans, multiple culture-based and culture-independent studies have identified the species as indigenous microbiota of various body sites, as reviewed by Scales, Dickson et al. (2014) who suggest a more complex relationship with human disease.

The Type VI secretion system (T6SS) is recently described in Gram-negative *Proteobacteriaceae* (Gerlach and Hensel 2007, Buchrieser, Rusniok et al. 2011). The T6SS system has been identified to be an important molecular mechanistic apparatus that is crucial for microbial interactions (Gerlach and Hensel 2007). There are 13 essential genes referred to as the core components of the T6SS that are conserved, however the order that the genes lie in differs in different species (Gerlach and Hensel 2007). Two of the core genes, *hcp* and *vgrG*, have been identified to code for the extracellular machinery and their proteins (they are predicted as structural and effector protein coding genes). They have been shown to be released if the T6SS system is functional (Pukatzki, McAuley et al. 2009). Many research studies have reported that the T6SS system is used by bacteria to kill competitive bacteria (Hood, Singh et al. 2010, MacIntyre, Miyata et al. 2010, Schwarz, West et al. 2010). Immune proteins are produced by bacteria to prevent self-intoxication by their own toxins. Such an interaction is observed in *Pseudomonas aeruginosa* Tsi proteins (immune proteins) and *Serratia marcescens* where Rap proteins (immune proteins) and the T6SS system deliver the toxins to the target cells to exert their antibacterial activity (Murdoch, Trunk et al. 2011, English, Trunk et al. 2012, Russell, Singh et al. 2012, Carruthers, Nicholson et al. 2013). Several *P. fluorescens* strains possess the genes encoding T6SS components and genomic and transcriptomic studies have suggested interactions with plants (Shrivastava and Mande 2008, Barret, Egan et al. 2011, Barret, Egan et al. 2013, Decoin, Barbey et al. 2014).

Considering both the harmful effects of chemical sanitizers and the potential of BCAs in controlling foodborne pathogens, we wanted to isolate bacteria with bio-preservative properties and understand their potential as BCA candidates. For this purpose, we divided the study in two parts. The first part involved isolation of resident bacterial species that have anti-bacterial activity (protective bacteria) from horticultural produce and/or their processing environments. The second part involved characterizing the species in terms of: (1) species identification by 16SrRNA sequencing; (2) analyzing the functional proteome using Liquid Chromatography with tandem-Mass-Spectrometry (LC-MS/MS); and (3) comparison of the T6SS secretion system among the strains that exhibited antibacterial activity as this is little studied.

## Materials and Methods

### Study part I

#### Screening for protective bacteria in horticultural produce

Resident bacteria with potential bio-control properties were isolated from apple samples (n=27) and swabs (n=7) from an apple pack house in Gisborne, New Zealand. The apples (n = 3 per bag of sample) were hand massaged in 400 mL of Butterfield broth (Difco - BD Becton Dickinson, and Company, Sparks, MD 21152, USA) and the swabs were stomached for 2 min in 100 mL Butterfield broth. The processed samples were incubated at 30°C for 24 h and 1-mL aliquots from the incubated broth samples were used to screen for resident bacteria with bio-preservative properties.

#### Pour plate screening

*L. monocytogenes* Scott A was grown in tryptic soy broth with 0.6% yeast extract (TSBYE) (Difco - BD, Becton, Dickinson and Company) for 24 h at 37°C and 1 mL. of the 24-h culture was used to make the *L. monocytogenes* pour plates using TSAYE (tryptic soy agar with yeast extract) (Difco - BD, Becton, Dickinson and Company). After solidification, 100 μL from the sample broths were spread using plate spreaders and left uncovered in a biosafety cabinet level II for 2 h to allow the samples to be absorbed. The spread pour plates were incubated at three different temperatures, 20, 30 and 37°C in an effort to provide good coverage to capture the resident bacteria with potential bio-control properties that grow at these temperatures.

#### Selection of colonies

Bacterial colonies that produced an identifiable zone of inhibition with Scott A were picked from the pour plates and regrown in TSBYE broth at the same respective temperatures. The cultures were then streaked onto TSAYE plates. If a mixture of cultures was found, single colonies of each morphology were streaked on *Listeria* CHROMagar (CHROMagar, 75006 Paris, France) to discriminate the *L. monocytogenes* colonies and non-*Listeria* cultures. The non-*Listeria* cultures were grown in TSBYE broth for 24 h at 20, 30 and 37°C and 100 μL spread onto *L. monocytogenes* Scott A pour plates to confirm the inhibitory activity of the respective colony cultures. The colonies that showed a zone of inhibition were further purified following the same procedure until a pure colony with a zone of inhibition was obtained. The pure colonies were propagated in Brain Heart Infusion agar (BHI, Difco - BD Becton, Dickinson and Company) plates and stored at −85°C in 50% glycerol broth until further testing.

#### Agar gel diffusion test for the zone of inhibition test

The purified presumptive bio-control cultures (n=14) that grew at 30°C (there was no bio-control property observed at other temperatures) and that showed some degree of inhibition were subjected to an agar gel diffusion test to confirm their antibacterial effect against *L. monocytogenes* Scott A and separately against other strains of *L. monocytogenes* from the Plant & Food Research culture collection (PFR18C07, PFR18D01 and PFR18D05). The *L. monocytogenes* strains were adjusted to an optical density (OD) of 0.5 at 600 nm and 1 mL of the culture was used in pour plates of TSAYE agar. Once solid, 4-mm diameter holes were aseptically punched into the agar with a sterile borer and the agar plugs were removed using sterile pipette tips. The presumptive cultures were grown in TSBYE broth for 24 h at 30°C and centrifuged at 4°C (Eppendorf, 5810R, Auckland, New Zealand) at 3,220 × *g* for 10 min. The supernatants were separated, filter sterilized using 0.2-micron syringe filters (Sartorius, Thermofisher Scientific) and stored at 4°C before further testing. The cell pellets were washed twice and resuspended with 0.1% tryptone (Difco - BD, Becton, Dickinson and Company), adjusted to an OD value of 0.5 at 600 nm and 30 μL added to holes in the pour plates. A chloramphenicol disc (30 μg, Mast Diagnostics, Mast group Ltd, Merseyside, United Kingdom) was placed in the center of each plate as a positive control. Plates were incubated at 30°C and 20°C and zones of inhibition recorded over a period of 5 days. The radius from the edge of the well to the edge of the clear zone was measured using a Vernier caliper (ROK Precision Instrument, Shenzhen, China). In the case of irregular edges, radii on all four sides of the inhibition zone were measured, and an average was calculated. Alternatively, the vegetative cells (10 μL) were placed on the pour plates to measure the zone of inhibition.

#### Testing of supernatant and vegetative culture of the isolates

Thirty μL of the sterile supernatant from all 14 cultures were dispensed into the wells (replicates of two plates were tested), left in the cabinet for 1 h for the liquid to be fully absorbed and then incubated at 20°C and 30°C for 24, 48 and 72 h. The OD values for the washed and resuspended cells were measured at 600 nm and adjusted to an OD of 0.5 for the agar diffusion test. Thirty μL of the re-suspended (0.5 OD) cells was dispensed into the wells and incubated at 30°C, 20°C and 37°C for 24, 48, 72 h and observed after 5, 10 and 15 days for any inhibition. For testing the technique of supernatants on pour plates, supernatants from protective cultures that have been proven to be listeriolytic *(Carnobacterium maltaromaticum* 1944, *C. maltaromaticum 2003, C. maltaromaticum, Carnobacterium divergens* 2122, *Leuconostoc gelidum* and *Lactococcus piscium)* were used as positive control supernatants in addition to antibiotic discs. The cultures were supplied by Institut Français de Recherche pour l’Exploitation de la Mer (IFREMER), Laboratoire de Génie Alimentaire, Nantes, France. Isolates that showed a minimum of a 1-mm zone of inhibition were chosen for species identification in Study part II: 16S rRNA gene analysis.

#### Species identification using 16SrRNA sequence analysis

The pure cultures were grown in TSAYE plates for 24 h and DNA was extracted using 2% Chelex solution (Chelex 100 resin, Bio-Rad laboratory, Hercules, California, USA). The colonies (2-3 mm) were picked using sterile loops, suspended in 2% Chelex solution and heat treated at 100°C for 10 min. The treated solution was cooled to room temperature, then centrifuged (Eppendorf, 5424R) at 21,130 × g for 5 min and 250 μL of the supernatant was separated and stored for PCR reactions. Isolates with DNA A_260_/ A_280_ ratios between 1.8 and 2.0 were taken for PCR amplification while DNA isolation was repeated for those not fulfilling this quality criterion until it was achieved. The DNA samples were amplified using Universal 16S rRNA bacterial primers 27F (5’-AGAGTTTGATCCTGGCTCAG-3’) and 1541R (5’-AAGGAGGTGATCCAGCCGCA-3’) (Zhou, Davey et al. 1997, Srinivasan, Karaoz et al. 2015).

The PCR reaction mix comprised of 10 μL of l× PCR buffer (Thermo Fisher Scientific), 2 μL of 10 μM DNTP mix (Thermo Fisher Scientific), 1 μL of 10 μM of each primer (forward and reverse, IDT, Australia), 1 μL Platinum Taq polymerase (Thermo Fisher Scientific) (one unit per reaction), MgCl_2_ 1.5 mM (Thermo Fisher Scientific) and 10 ng/μL of DNA. The reaction mix was made up to 50 μL with sterile MilliQ water. The PCR reaction was carried out in an Eppendorf Master Gradient Cycler with the following conditions: initial denaturation at 95°C for 2 min, 95°C for 30 sec, annealing at 55°C for 30 sec, elongation at 72°C for 2 min with a further elongation of 10 min. The reaction was carried out for 35 cycles.

PCR products were viewed under 1% agarose gel (Ultrapure™ Agarose, Invitrogen, Thermo Fisher Scientific, Massachusetts, USA) stained with RedSafe Nucleic Acid Staining Solution (JH Science / iNtRON Biotechnology USA) using a Gel Documentation system (BioRad, Hercules, California, USA) to confirm the presence the PCR product (approximately 1,464 bp). PCR products were purified using a QIAquick PCR Purification Kit (Qiagen) and the products were quantified using Nanodrop and sent to Macrogen Inc, Korea at a concentration of 20 μg/μL for sequencing. The Fasta sequences of all the cultures were blasted using NCBI BLAST blasting suite (https://blast.ncbi.nlm.nih.gov/Blast.cgi). The sequences were identified to genus level where an unambiguous high identity and coverage of ≥ 95 to 98% were taken as the criteria for determining the genus of the DNA sequences (Barghouthi 2011). The sequence analyses of the isolates were carried out in Geneious v10.1 and a Neighbor-Joining phylogenetic tree was constructed in MEGA v7, using the Bootstrap method (1000 replications). The forward and reverse sequences were compared individually, and a consensus was used for the comparison and phylogenetic tree construction. In addition to 16SrRNA identification, the proteomic data analysis described in the following section confirmed the species.

#### Testing of cultures for fluorescence activity and zone of inhibition

Isolates that were 16SrRNA sequenced and identified as *Pseudomonas fluorescens* were subsequently tested for fluorescence activity by growth on BHI agar plates, exposure to UV irradiation (BioRad) and image capture. TSAYE pour plates were prepared from 35 randomly selected strains (Table 1) of *L. monocytogenes* sourced from seafood (n=17), horticultural produce (n=14), clinical isolates (n=2) plus one *L. innocua* and one *L. seeligeri*. A positive control chloramphenicol disc (30 μg,) was used in the center of the plate for each strain. *L. innocua* and *L. seeligeri* were used in the trial to find out if the organisms are capable of inhibiting other *Listeria* spp. as well.

**Table 1:** *Listeria monocytogenes, L. seeligeri* and *L. innocua* strains used in the study and their New Zealand sources

| No. | Strain | Organism | Source |
| --- | --- | --- | --- |
| 1 | PFR05A12 | *L. seeligeri* | Vegetable |
| 2 | PFR12C05 | *L. monocytogenes* Scott A | Clinical isolate |
| 3 | PFR16B03 | *L. monocytogenes* ATCC strain 49594 | Clinical isolate |
| 4 | PFR18B09 | *L. monocytogenes* | Seafood processing environment |
| 5 | PFR18C05 | *L. monocytogenes* | Seafood processing environment |
| 6 | PFR18C07 | *L. monocytogenes* | Seafood processing environment |
| 7 | PFR18D01 | *L. monocytogenes* | Seafood processing environment |
| 8 | PFR18D05 | *L. monocytogenes* | Seafood processing environment |
| 9 | PFR33F02 | *L. monocytogenes* | Seafood processing environment |
| 10 | PFR33F03 | *L. monocytogenes* | Seafood processing environment |
| 11 | PFR33H03 | *L. monocytogenes* | Seafood processing environment |
| 12 | PFR33H04 | *L. monocytogenes* | Seafood processing environment |
| 13 | PFR33I04 | *L. monocytogenes* | Seafood processing environment |
| 14 | PFR40I05 | *L. monocytogenes* | Horticultural source |
| 15 | PFR40I07 | *L. monocytogenes* | Horticultural environment |
| 16 | PFR41E01 | *L. monocytogenes* | Horticultural environment |
| 17 | PFR41E02 | *L. monocytogenes* | Horticultural environment |
| 18 | PFR41E03 | *L. monocytogenes* | Horticultural environment |
| 19 | PFR41E05 | *L. monocytogenes* | Horticultural environment |
| 20 | PFR41F08 | *L. monocytogenes* | Horticultural environment |
| 21 | PFR41G01 | *L. monocytogenes* | Horticultural environment |
| 22 | PFR41G02 | *L. monocytogenes* | Horticultural environment |
| 23 | PFR41H07 | *L. monocytogenes* | Horticultural environment |
| 24 | PFR41J05 | *L. monocytogenes* | Horticultural environment |
| 25 | PFR41J08 | *L. monocytogenes* | Horticultural environment |
| 26 | PFR41J09 | *L. monocytogenes* | Horticultural environment |
| 27 | PFR42G03 | *L. monocytogenes* | Horticultural environment |
| 28 | PFR42I05 | *L. monocytogenes* | Seafood processing environment |
| 29 | PFR42I06 | *L. monocytogenes* | Seafood processing environment |
| 30 | PFR42I07 | *L. monocytogenes* | Seafood processing environment |
| 31 | PFR42I08 | *L. monocytogenes* | Seafood processing environment |
| 32 | PFR42I09 | *L. monocytogenes* | Seafood processing environment |
| 33 | PFR42I10 | *L. monocytogenes* | Seafood processing environment |
| 34 | PFR42J03 | *L. monocytogenes* | Seafood processing environment |
| 35 | PFR05A10 | *L. innocua* | Processed vegetable |

#### Testing for antibacterial activity of *P. fluorescens* in liquid media in co-culture

Cultures of *L. monocytogenes* strains were adjusted to ODs of 0.5 at 600 nm as were *P. fluorescens* isolates. Both microbes were co-cultured in TSBYE by adding 50 μL of each strain of *P. fluorescens* individually with each *Listeria* strain and serial dilution to 10^−7^. The co-culture was incubated at 30°C, 20°C and 37°C for 24, 48, 72 h and observed after 5, 10 and 15 days. After each time point, 10 μL of the co-culture from all dilutions was plated onto ChromAgar *Listeria* and TSAYE plates to observe the growth of blue colonies for *Listeria* spp. on ChromAgar and white or creamy colonies on TSAYE for *P. fluorescens*.

#### Proteome analysis of *P. fluorescens*

The pure cultures identified as *P. fluorescens* using 16S rRNA were subjected to proteomic analysis using whole cell lysates. Isolate PFR46I06 did not exhibit fluorescence and did not have profound zone of inhibition so only PFR46H06, PFR46H07, PFR46H08 and PFR46H09 were subjected to proteome analysis. The cultures were grown in BHI broth for 24 h and were centrifuged at 3200 × g for 20 min at 4°C (Eppendorf, 5424R). The cell pellets were transported on ice for proteome analysis at The Mass Spectrometry Centre, Faculty of Sciences, University of Auckland using a nanoLC-equipped TripleTOF 6600 mass spectrometer (ABSCIEX, USA) and utilizing Information Dependent Acquisition (IDA) method. The sample process involved cysteine alkylation using iodoacetamide and the samples were digested with trypsin with urea denaturation. The ProteinPilot data were searched using UniProt protein database of *Pseudomonas fluorescens* sequences (October 2018). The summaries of proteins, peptides and distinct peptides with modifications were analysed with special reference to the secretion system, T6SS. The mass spectrometry proteomics data have been deposited to the ProteomeXchange Consortium via the PRIDE [1] partner repository with the dataset identifier PXD019965.

#### Statistical analysis

Statistical analyses were carried out using R version 3.5.1. Each experiment was carried out twice, for agar gel diffusion tests and for the co-culture methods, and mean, standard deviation and standard errors were calculated and one way and two way ANOVAs and post-hoc Fisher’s least significant difference (LSD) Bonferroni with an alpha error of 0.05 were calculated using the Agricolae ™ package in R for each strain of *P. fluorescens* against 35 strains of *Listeria* spp. A logistic regression model was used to compare the *L. monocytogenes* strains and the zone of inhibition produced by each strain of *P. fluorescens*. This model consisted of zones of inhibition produced by the three *Pseudomonas* strains that produced recognizable zones of inhibition on all *Listeria* strains tested. The model had the *Listeria* strains and *Pseudomonas* strains as influencing variables. (*Listeria* strains vs PFR46H07+ PFR46H08+ PFR46H09).

## Results

### Bacterial cultures screening, 16SrRNA identification and fluorescence testing

Of the 27 apple and seven environmental swab samples screened for resident bacteria with bio-control characteristics, five cultures (PFR46H06, PFR46H07, PFR46H08, PFR46H09 and PFR46I06) were isolated with small scale (maximum of 2 mm zone of inhibition) to identifiable scale (3-5 mm) of listericidal activity. Of the five isolates, three (PFR46H07, PFR46H08 and PFR46H09) exhibited recognizable zones of inhibition (larger than 2 mm) while the other two showed less than 1.5 mm zones of inhibition. The five cultures were subjected to 16S rRNA gene sequencing for species identification and identified as *P. fluorescens*, based on the sequence identity score (above 95%). Figure 1 shows the Neighbor-Joining phylogenetic tree of the five isolates along with best matching blast sequences that had above 95% sequence identity matching. Four of these five cultures when grown on BHI agar plates had a light greenish color and fluoresced under UV light (Supplementary Figure 1a), while PFR46I06 did not show bright fluorescence (Supplementary Figure 1b). The proteome from all isolates matched *P. fluorescens* and species identity was confirmed.

**Figure 1:**
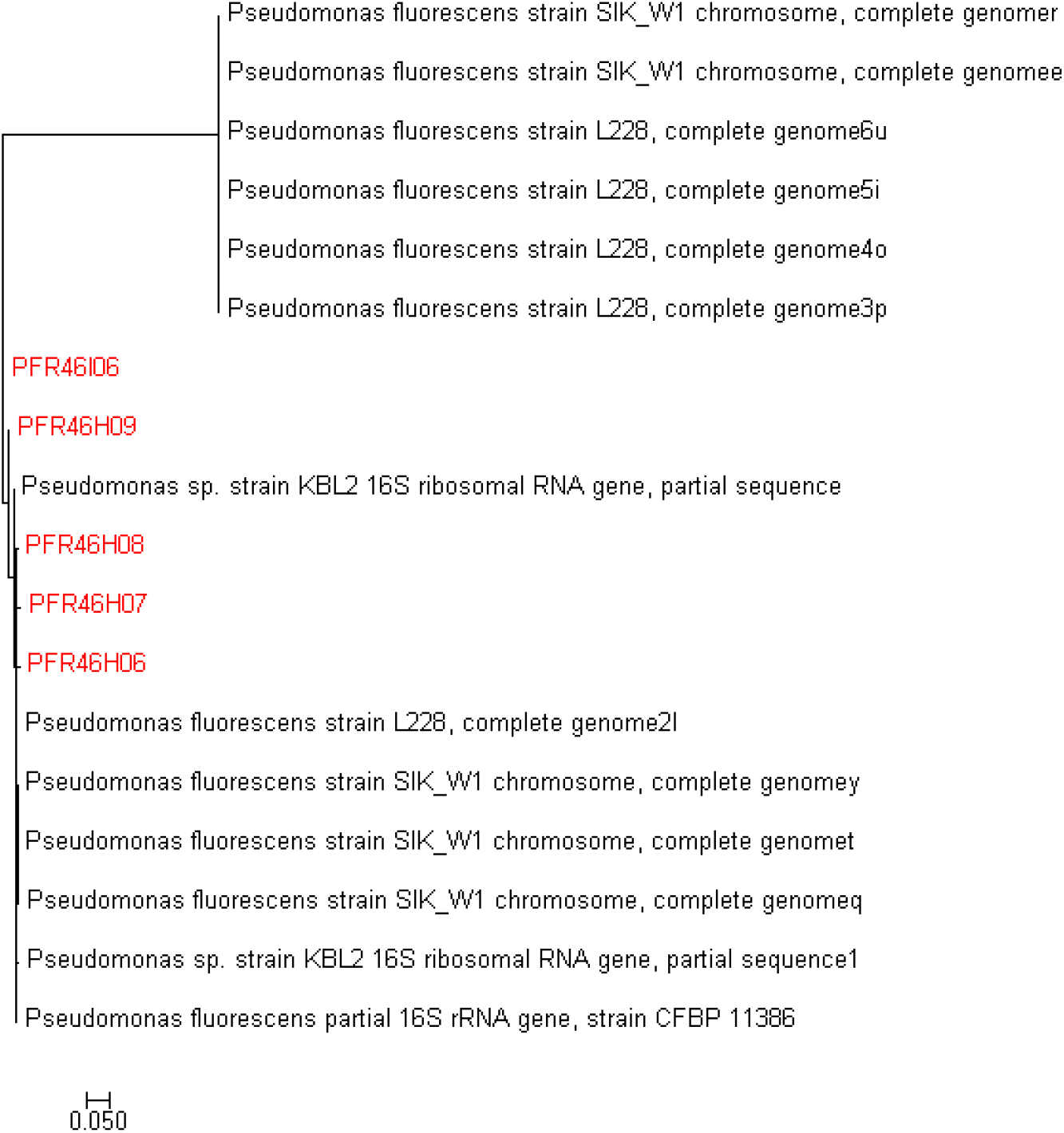
Neighbor-Joining Tree of the 16SrRNA sequences of 13 reference *Pseudomonas fluorescens* isolates that had a sequence identity above 95% in blast analysis along with the five isolates of *P. fluorescens* isolated in this study constructed using MEGA v7 with Bootstrap value of 1000 replications. The forward and the reverse sequences were compared individually, and a consensus was used for the comparison and phylogenetic tree construction.

### Zone of inhibition

Three fluorescent isolates (PFR46H07, PFR46H08 and PFR46H09) showed a detectable zones of inhibition at both 20 and 30°C, however, the zones of inhibition were clearer and more defined on the culture plates at 20°C after 48 h (Supplementary Figure 2) than 24 hours. The isolates PFR46H06 and PFR46I06 showed very small zones of inhibition towards some strains that could not be measured (less than 1.5 mm) and were not inhibitory to the majority of the *L. monocytogenes* isolates. Therefore, the three strains that showed a minimum zone of inhibition of ca 10 mm and above were recorded for the purpose of comparison and interpretation. The average inhibition zone for the culture plates grown at 30°C was 10 mm, 9.5 mm and 10.5 mm collectively for all *Listeria* strains for PFR46H07, PFR46H08 and PFR46H09, respectively. One way ANOVA comparing individual strains against 35 *L. monocytogenes* strains showed no significant difference in the zone of inhibition between the three strains. Figure 2 shows the inhibition zones produced by the three strains against 35 *Listeria* strains. One way ANOVA individually comparing the zone of inhibition variances between the three *P. fluorescens* strains showed no significant differences against the *Listeria* spp. compared (P ≥ 1). However, the logistic regression model showed that the intercept was highly significant (P < 0.00), and the inhibition zones produced towards the *L. monocytogenes* strains PFR18C07 and PFR18D05 were significantly different compared with the other *Listeria* strains (Table 2). In contrast, the supernatants did not show any inhibition.

**Table 2:** Logistic regression model of the inhibition zones produced by three *Pseudomonas fluorescens* strains against 33 *Listeria monocytogenes,* one *Listeria innocua* (PFR05A10) and one *Listeria seeligeri* (PFR05A12) strains collected from New Zealand seafood, seafood processing environments, and horticultural sources (PFR05A10 has been taken as a reference in the model by default). Bold fonts represent P values being either, highly significant (underlined, P<0.05, ***), significant (underlined, P<0.05, *) or borderline (P≥0.05 and P≤0.10,.)

| *Listeria* strains |  |  |  |  |
| --- | --- | --- | --- | --- |
|  | Estimate | Std. Error | z value | Pr(>\|z\|) |
| PFR05A10 |  |  |  |  |
| (Intercept) | 2.32 | 0.58 | 4.00 | **<u>0.00</u>** *** |
| PFR05A12 | 0.13 | 0.78 | 0.17 | 0.86 |
| PFR12C05 | -0.18 | 0.73 | -0.25 | 0.80 |
| PFR16B03 | -0.34 | 0.70 | -0.48 | 0.63 |
| PFR18B09 | -1.06 | 0.64 | -1.66 | **0.10** . |
| PFR18C05 | -1.13 | 0.63 | -1.80 | **0.07** . |
| **<u>PFR18C07</u>** | -1.48 | 0.61 | -2.43 | **<u>0.02</u>** * |
| PFR18D01 | -1.16 | 0.63 | -1.85 | **0.06** . |
| **<u>PFR18D05</u>** | -1.41 | 0.62 | -2.28 | **<u>0.02</u>** * |
| PFR33F02 | -0.49 | 0.69 | -0.71 | 0.48 |
| PFR33F03 | -0.37 | 0.70 | -0.52 | 0.60 |
| PFR33G10 | -0.47 | 0.69 | -0.68 | 0.50 |
| PFR33H03 | -0.43 | 0.70 | -0.62 | 0.54 |
| PFR33H04 | -0.44 | 0.70 | -0.64 | 0.53 |
| PFR40I05 | -0.83 | 0.67 | -1.23 | 0.22 |
| PFR40I07 | -0.28 | 0.72 | -0.40 | 0.69 |
| PFR41E01 | 0.05 | 0.76 | 0.07 | 0.95 |
| PFR41E02 | -0.13 | 0.73 | -0.18 | 0.86 |
| PFR41E03 | -0.62 | 0.68 | -0.91 | 0.37 |
| PFR41E05 | -0.91 | 0.66 | -1.38 | 0.17 |
| PFR41F08 | -0.35 | 0.71 | -0.49 | 0.62 |
| PFR41G01 | 0.01 | 0.75 | 0.01 | 0.99 |
| PFR41G02 | 0.01 | 0.75 | 0.01 | 0.99 |
| PFR41H08 | -0.08 | 0.74 | -0.11 | 0.91 |
| PFR41J05 | -0.17 | 0.72 | -0.24 | 0.81 |
| PFR41J08 | -1.17 | 0.63 | -1.87 | **0.06** . |
| PFR41J09 | -0.23 | 0.72 | -0.31 | 0.75 |
| PFR42G03 | -0.36 | 0.70 | -0.51 | 0.61 |
| PFR42I05 | -0.43 | 0.70 | -0.61 | 0.54 |
| PFR42I06 | -0.32 | 0.71 | -0.46 | 0.65 |
| PFR42I07 | -0.26 | 0.71 | -0.37 | 0.72 |
| PFR42I08 | -0.31 | 0.71 | -0.43 | 0.67 |
| PFR42I09 | -0.26 | 0.71 | -0.36 | 0.72 |
| PFR42I10 | -0.33 | 0.71 | -0.46 | 0.65 |
| PFR42J03 | -0.29 | 0.71 | -0.41 | 0.68 |

**Figure 2:**
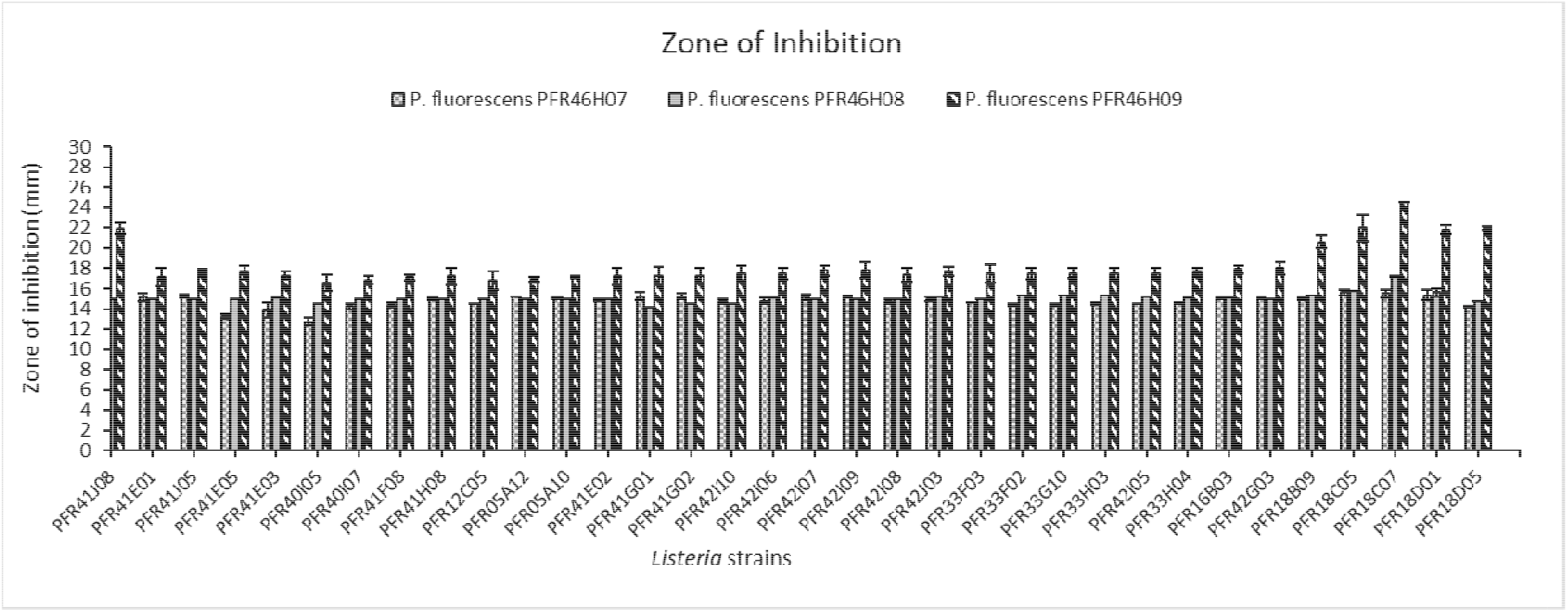
Inhibition zones (two replicates averaged) produced by three *Pseudomonas fluorescens* strains against 31 *Listeria monocytogenes*, one *Listeria innocua* (PFR05A10) and one *Listeria seeligeri* (PFR05A12) strains collected from seafood and horticultural sources in New Zealand and two international clinical isolates (*L. monocytogenes* ScottA = PFE12C05 and ATCC 49594 = PFE16B03).

### Testing for antibacterial activity in liquid media in co-culture

There was no evidence of inhibition of *Listeria spp.* when co-cultured with any of the *P. fluorescens* isolates in liquid media. The cultures were very slimy and ropy and difficult to pipette or plate as the number of days in co-culture increased. In contrast, the inhibition in solid media increased as the number of days in culture increased.

### Proteome analysis of *P. fluorescens* strains

We analyzed the proteomes of the four fluorescing isolates, one that did not have profound inhibition (PFR46H06) and three inhibitory isolates (PFR46H07, PFR46H08 and PFR46H09). Figure 3 shows the false discovery rates of proteins at 1%, 5% and 10% error rates compared with global protein databases. The proteome analysis further confirmed the isolates as *P. fluorescens*. All four isolates showed different protein hits with 1781 in PFR46H06, 2030 in PFR46H07, 2228 in PFR46H08 and 1994 in PFR46H09. The lowest number of proteins (1994) was in PFR46H09, the strain with the strongest antimicrobial properties, suggesting its proteome may be small compared with the other three isolates. However, due to the liquid co-culture results, our main interest was in investigating the secretion systems with special reference to the T6SS in each of the isolates.

**Figure 3:**
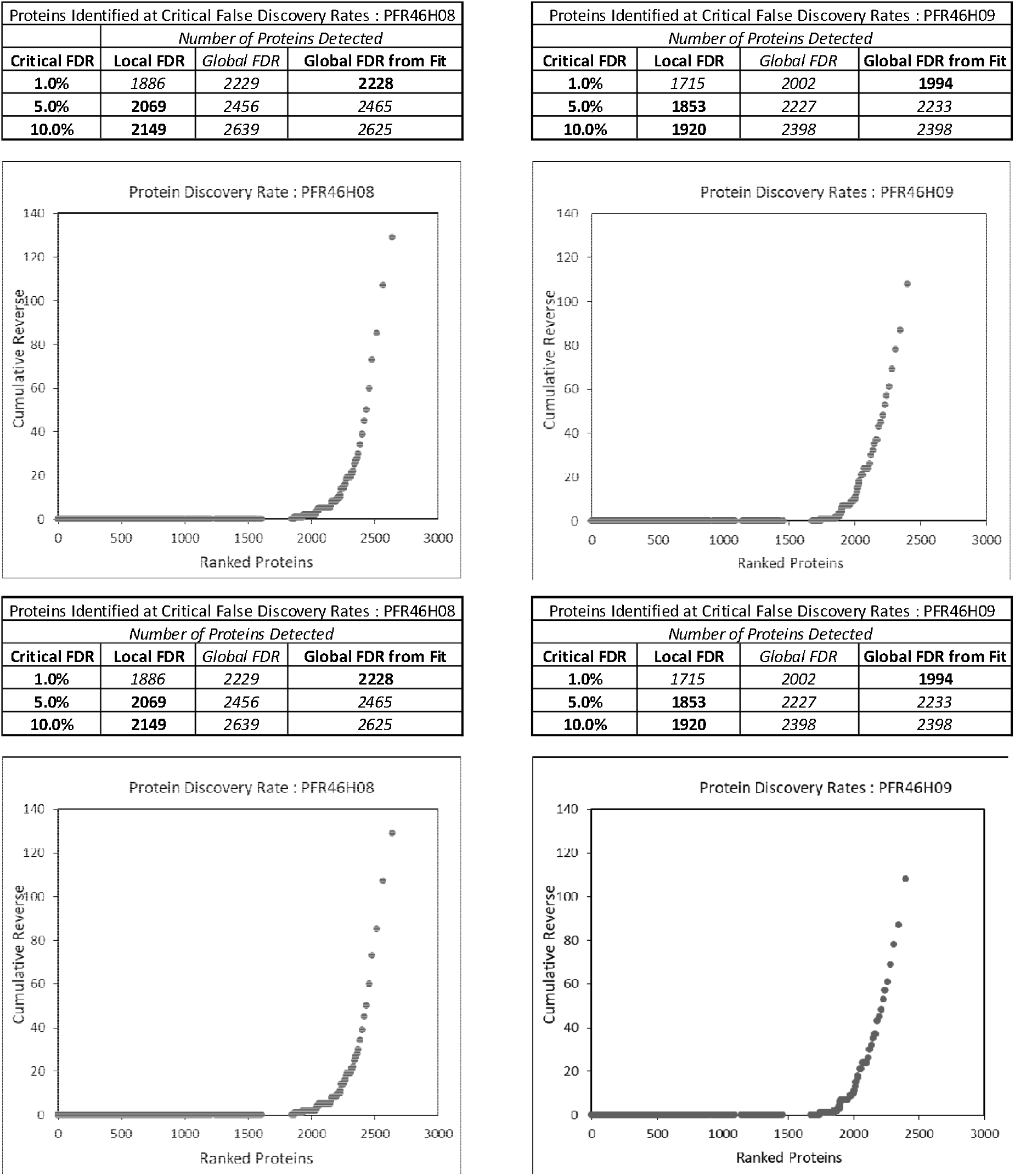
The number of proteins at false discovery rates (FDR) of 1, 5 and 10% for the *Pseudomonas fluorescens* isolates. The total number of proteins varied in each isolate with PFR46H06, PFR46H07, PFR46H08 and PFR46H09 showing 1781, 2030, 2228 and 1994 proteins, respectively.

We compared the different secretion systems, including fimbria and flagella related proteins, phage and phage related proteins and hemolysin proteins, and found substantial differences in the number of proteins among the four isolates. Supplementary Table 1 lists all the proteins that were detected in the proteomes of each *P. fluorescens* isolate while Table 3 lists the secretion system proteins. PFR46H06 and PFR46H07 possessed the fewest secretion proteins (12 and 11 respectively) while PFR46H08 and PFR46H09 each had 18. PFR46H09, which showed the greatest antimicrobial effect, had nine T6SS proteins compared to just four in the other three strains.

**Table 3:**
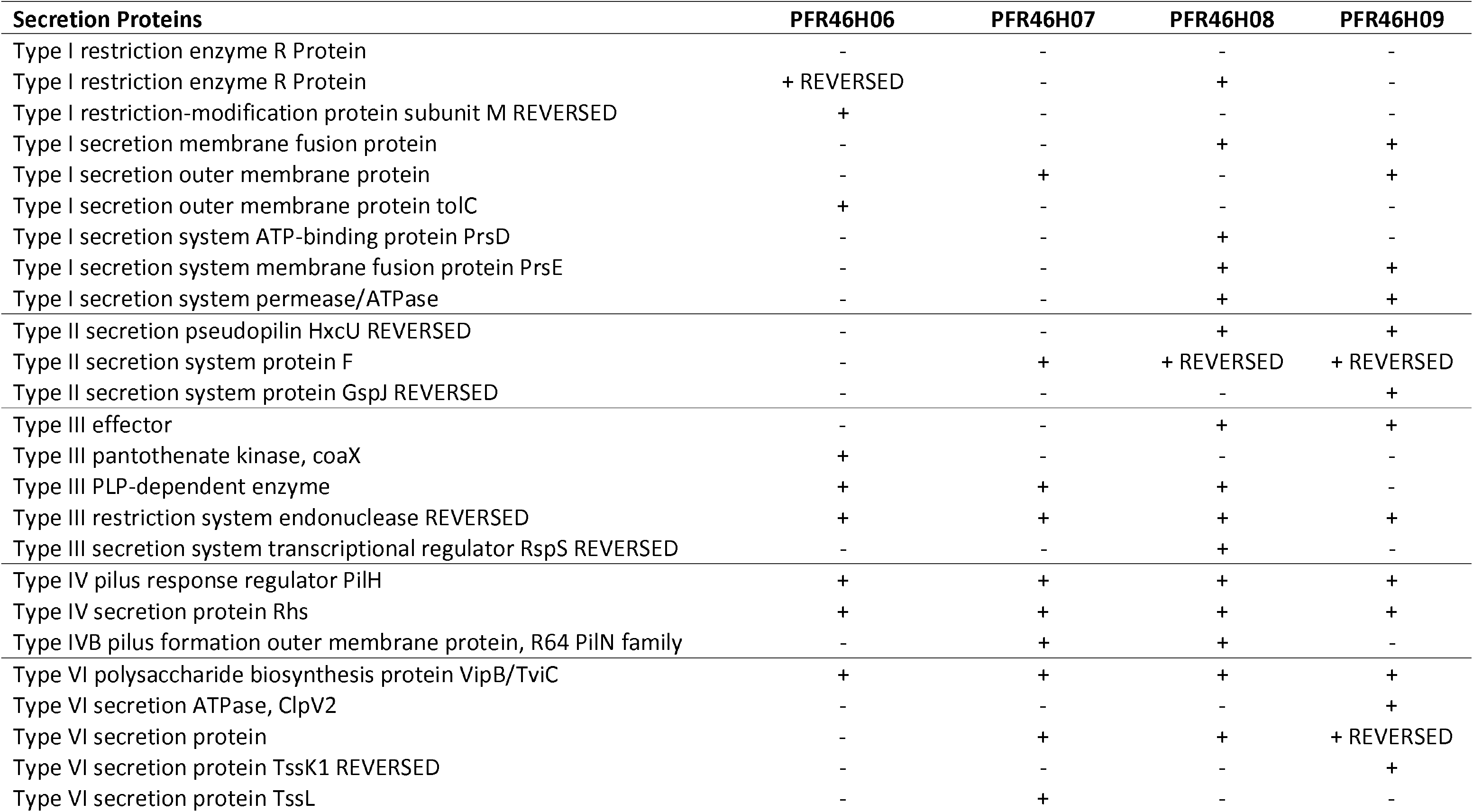
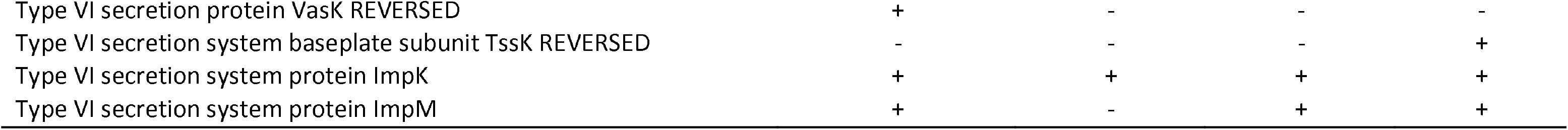
Summary of the different secretion system proteins detected in four *Pseudomonas fluorescens* proteomes: PFR46H06, PFR46H07, PFR46H08 and PFR46H09

T6SS protein ImpK was present in all four isolates. However, there were notable modifications in the protein. In PFR46H06 and PFR46H09, a protein modification, substituting from the amino acid R to N, was found at the 95^th^ position, while in PFR46H07 this R to N substitution occurred at the 92^nd^ position. In PFR46H08, the ImpK protein had a number of substitutions and modifications (Supplementary excel data sheets for individual strains with distinct peptide summaries): at 63, A to G; at 64, N to M; at 67, V to M; at 68, E to D, at 70, V to M; and at 95, R to N.

The next protein that we observed closely was the *rhs* gene protein of the T6SS. Protein modifications were found at the positions 37 (T to V) and 427 (S to T) in PFR46H06, PFR46H07 and PFR46H08 while in PFR46H09 there was an additional modification found at the 427^th^ position (S to T). Similarly, protein modifications were found in other secretion proteins in the T6SS system in PFR46H07, PFR46H08 and PFR46H09 such as having a Type VI secretion protein similar to the Type VI secretion system contractile sheath small subunit *VipA* in other *Pseudomonas* spp. (Kudryashev, Wang et al. 2015, Gallique, Bouteiller et al. 2017). In general, the number of secretion proteins found in PFR46H08 and PFR46H09 was greater compared with the other two isolates. The T6SS system proteins ClpV2, TssK1, TssK, VipB, and ImpM were present in PFR46H09 while the other isolates lacked one of more of these proteins (Table 3).

## Discussion

In the present study we isolated five strains of *P. fluorescens*, of which three showed significant anti-listerial activity in solid media, inhibiting all 35 strains of *Listeria* spp. tested. The zones of inhibition were more pronounced at 20°C than 30°C where the zones were hazy and not very clear (Supplementary Figure 2). PFR46H09 strain was significantly (P = 0.02) more inhibitory than the other two strains (PFR18D05 and PFR18C07, Table 2).

*Pseudomonas* is a noted genus of psychrotrophic spoilage organism found in soil, water, and vegetation (Palleroni 1984, de Oliveira, Favarin et al. 2015, Ribeiro Júnior, de Oliveira et al. 2018). Pseudomonads are also commonly found in unpasteurized milk and dairy products (de Oliveira, Favarin et al. 2015). Although Pseudomonads are reported to enhance growth of non-pathogenic and pathogenic bacteria in dairy products (Miller, Scanlan et al. 1973, de Oliveira, Favarin et al. 2015, Ribeiro Júnior, de Oliveira et al. 2018), studies also suggest that *P. fluorescens* has antibacterial activity against certain foodborne pathogens such as *L. monocytogenes* (Farrag and Marth 1989, Cheng, Michael et al. 1995, Hoffland, Hakulinen et al. 1996). *P. fluorescens* has therefore been proposed as a BCA (Weller 2007) and fluorescent Pseudomonads have been studied for biocontrol research since 1970 when the process was known as bacterization (Kloepper, Lifshitz et al. 1988). Farrag and Marth (1989) found that *P. fluorescens* moderately inhibited *L. monocytogenes* in skim milk stored at 7 and 13°C but that there was an enhancement of growth by *L. monocytogenes* Scott A in the presence of *P. fluorescens* P26 after 7 days of incubation at 7°C. A similar observation was made by Douglas and Schimdt (1988) at 10°C, however, in this trial, after 14 days the populations declined compared with controls. Both studies suggested a limited inhibitory effect of *P. fluorescens* against *L. monocytogenes* which was evident at low temperatures. In our observations, we show that there are differences among *P. fluorescens* strains where only three of five strains caused inhibition and different *L. monocytogenes* strains had different responses towards these strains.

We conducted proteomic analyses of the four strains of *P. fluorescens* with differences in zones of inhibition to help understand the potential of this bacteria to be a broad range BCA. Paul, Dineshkumar et al. (2006) previously examined the proteome of *P. fluorescens* MSP-393 in an effort to investigate the osmotolerance and/or saline stress levels of this strain to be used in agriculture production. Similarly, Kim, Silby et al. (2009), examined the proteome and identified the non-annotated protein coding genes in *P. fluorescens* strain Pf01. In our study, we carried out whole cell lysate proteome analysis and looked at the secretion systems, fimbria and flagella related proteins, mainly concentrating on the T6SS secretion system to see the differences between these strains. The T6SS system is one such newly identified secretion system (Hood, Singh et al. 2010, Gallique, Bouteiller et al. 2017).

In general, Gram-negative bacteria have been shown to utilize various secretion systems to deliver molecules into other bacterial and/or target cells as well as extracellular surfaces and these systems are considered to be important virulence factors, as reviewed by Costa, Felisberto-Rodrigues et al. (2015). Studies of the T6SS system indicate that approximately 15 conserved and closely linked genes are necessary to form a functional apparatus (Filloux 2009), and this apparatus is required to transport the hemolysin co-regulated protein and the valine-glycine repeat (Vgr) family proteins (Mougous, Cuff et al. 2006). Recent X-ray crystallographic studies (Leiman, Basler et al. 2009, Pell, Kanelis et al. 2009) suggested that these proteins are similar to bacteriophage tube and tail-spike proteins and speculated that T6SS could be evolutionarily, structurally, and mechanistically related to bacteriophage, which is bactericidal. The fundamental understanding of the mode of action of the T6SS changed significantly after the discovery that numerous critical components of T6SS are functionally homologous to the structural components of contractile phage tails (Costa, Felisberto-Rodrigues et al. 2015). The Hcp protein group was shown to be a structural homolog of phage tube proteins. In *Pseudomonas aeruginosa*, Hcp1 was shown to be the most abundant T6SS secreted protein and structurally has been shown to be a donut-shaped hexamer (Mougous, Cuff et al. 2006). These hexamers were shown to stack on top of each other head-to-tail to form continuous tubes in crystals that are identical to the external and internal proteins of the bacteriophage T4 tail tube (Ballister, Lai et al. 2008). In our proteomic study we observed several phage related tube, tail and sheath proteins to be present in all four strains, however, each strain was different in its respective protein summary. The strains of *P. fluorescens* had proteins that are either T7 tail tube proteins or homologs of T7 tail, tube and sheath associated proteins.

We observed that PFR46H09 possessed fewer total proteins but exhibited relatively larger inhibition zones compared with other strains. This strain also possessed relatively more T6SS proteins and flagella and phage related structural proteins. PFR46H09 colonies were irregular in shape and the presence of numerous flagella related proteins explains the movement on the solid agar media compared with other strains. Another essential conserved T6SS protein, TssE, has been shown to be homologous to T4 phage baseplate (Nano and Schmerk 2007, Leiman, Basler et al. 2009, Lossi, Dajani et al. 2011). In our study there were baseplate proteins: TssL in PFR46H07 and Tssk and TssK1 proteins in PFR46H09 (Table 3). Bonemann, Pietrosiuk et al. (2009) showed that VipA (TssB) and VipB proteins form the tubular polymer and Leiman, Basler et al. (2009) showed the overall structure resembled the T4 phage poly-sheath and can be disassembled by ClpV substrate protein. In our study, PFR46H09 revealed ClpV2 and all four strains showed VipB proteins (Supplementary Table 2).

Similarly, another study of T6SS in *Vibrio cholerae* established a model of the T6SS structure and its mode of action (Basler, Pilhofer et al. 2012). T6SS was shown to resemble a long phage tail that is attached to a cell envelope through an anchor. The tail was shown to have two conformations, extended and contracted, and this resembled the VipA/VipB sheath (Bonemann, Pietrosiuk et al. 2009, Basler, Pilhofer et al. 2012). The contracted sheath structures were in general shorter, wider and the T6SS sheath assembly in *V. cholerae* was shown to take about 20-30 s and then the sheath contracted to about half its length in less than 5 ms. This sheath was shown to disassemble in the presence of ClpV (Basler, Pilhofer et al. 2012). The T6SS dynamics were studied in a detailed manner using live-cell imaging in *V. cholerae, P. aeruginosa* and *Escherichia coli* (Basler, Pilhofer et al. 2012, Brunet, Espinosa et al. 2013, Kapitein, Bonemann et al. 2013). In summary, the roles of the T6SS conserved proteins are still deemed to be largely unknown (Hood, Singh et al. 2010) although T6SS has been shown to have a different mode of action from other secretion systems (Basler, Pilhofer et al. 2012, Brunet, Espinosa et al. 2013, Kapitein, Bonemann et al. 2013).

T6SS protein ImpK was found to be present in all four of our isolates and this protein is thought to be similar to *E. coli* outer-membrane protein, and to the flagellar torque-generating protein that was discovered by a study in 2003 (Bladergroen, Badelt et al. 2003). There were several notable differences in the protein modifications in all four isolates. In PFR46H06 and PFR46H09, a protein modification was found at the 95th position, from R to N, while in PFR46H07 the substitution had occurred at the 92^nd^ position and in PFR46H08, quite a number of substitutions and modifications were detected compared with the global *P. fluorescens* protein databases (Supplementary Table 2).

The rhs protein family is known to be widely spread in Gram-negative bacteria (Kung, Khare et al. 2012, Jones, Hachani et al. 2014, Alteri and Mobley 2016). All of our strains possessed the *rhs* gene of the T6SS, with modifications in some strains compared with the reference strains in the UniprotKB database. PFR46H09 possessed a protein modification at the 427^th^ position from S to T which made this protein different from the other three strains. Similarly, protein modifications were found in the T6SS contractile sheath small subunit *VipA* that had been identified in previous studies (Kudryashev, Wang et al. 2015, Gallique, Bouteiller et al. 2017).

We found that Imp family proteins were also present in all four isolates. A 2003 study reported a putative operon of 14 genes in *Rhizobium leguminosarum* strain RBL5523 and named them impA-impN (Bladergroen, Badelt et al. 2003). ImpK (present in all strains) resembles an outer membrane protein gene of *E. coli* and the authors suggested that the Imp system encoded components of a secretion apparatus and the proteins dependent on Imp genes blocked colonization/infection processes in pea plants (Bladergroen, Badelt et al. 2003).

PFR46H06, which had minimal inhibition of *Listeria*, lacked two secretion proteins that were present in the other three inhibitory strains (Type II secretion system protein F and Type VI secretion protein). This may explain its poor or no inhibition. Among the T6SS proteins, PFR46H06 did possess a transposon mediated virulence protein vasK which was not found in the other isolates and this protein has been proposed be associated with the T6SS system that is required for cytotoxicity of *V. cholera* cells toward *Dictyostelium* amoebae (Pukatzki, Ma et al. 2006).

We acknowledge that these strains were cultured in TSBYE without aiming for conditional expressions of the genes. Therefore, these strains may exhibit different characteristics at least in terms of inhibition of other bacteria under different environmental conditions and their proteomes may differ under respective conditions. We observed little or no inhibition of *Listeria* in liquid media. This was also observed by Silverman with *P. aeruginosa* poorly expressing the T6SS system in liquid medium, while it was expressed during surface growth (Silverman, Austin et al. 2011).

In conclusion, our study has evaluated the listeriolytic activity of four strains of *P. fluorescens* isolated from horticultural environments through zone of inhibition testing, identifying three strains with strong inhibition and one strain that produced minimal listeriolytic effect. We have identified some probable mechanisms in the antibacterial effects of the *P. fluorescens* isolates through proteomic analysis and particularly by concentrating on the T6SS system in each strain. However, the machinery of the T6SS system in *P. fluorescens* needs further investigation in order to better understand the exact mechanisms of action and their potential benefits to be used as an efficient antibacterial and/or listeriolytic agent.

## Supporting information

https://drive.google.com/drive/folders/1ZlbR6UaR0kC5LnZymSXj3ZuSl8qdavyN

## Acknowledgements

We thank Reginald Wibisono and Saili Chalke from Plant & Food Research for their help in this project. We also acknowledge funding from Plant & Food Research’s Discovery Science and Strategic Science Investment Funds, Future Consumer Foods Programme 1921.

---

**Supplementary Table 1:**
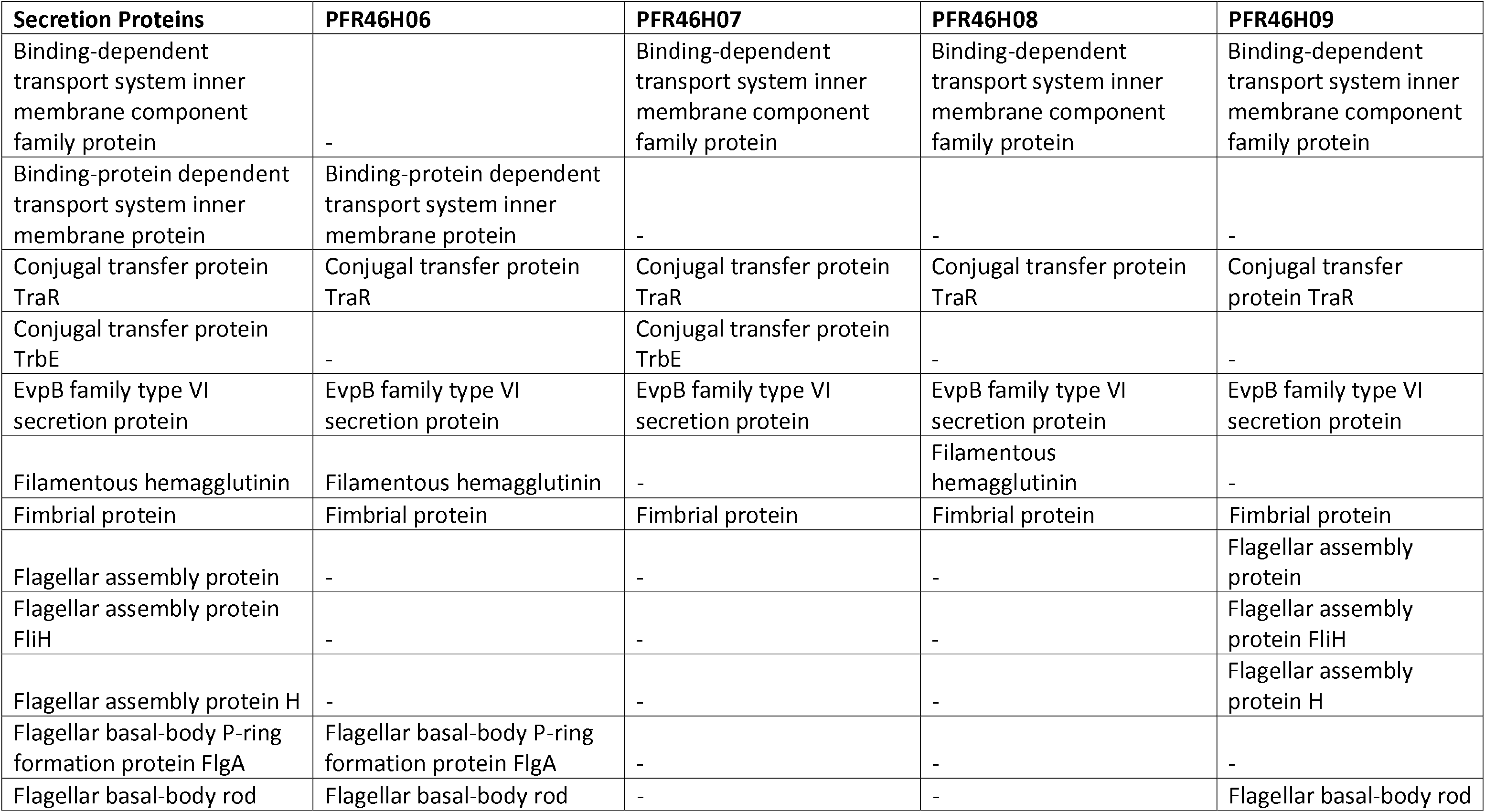
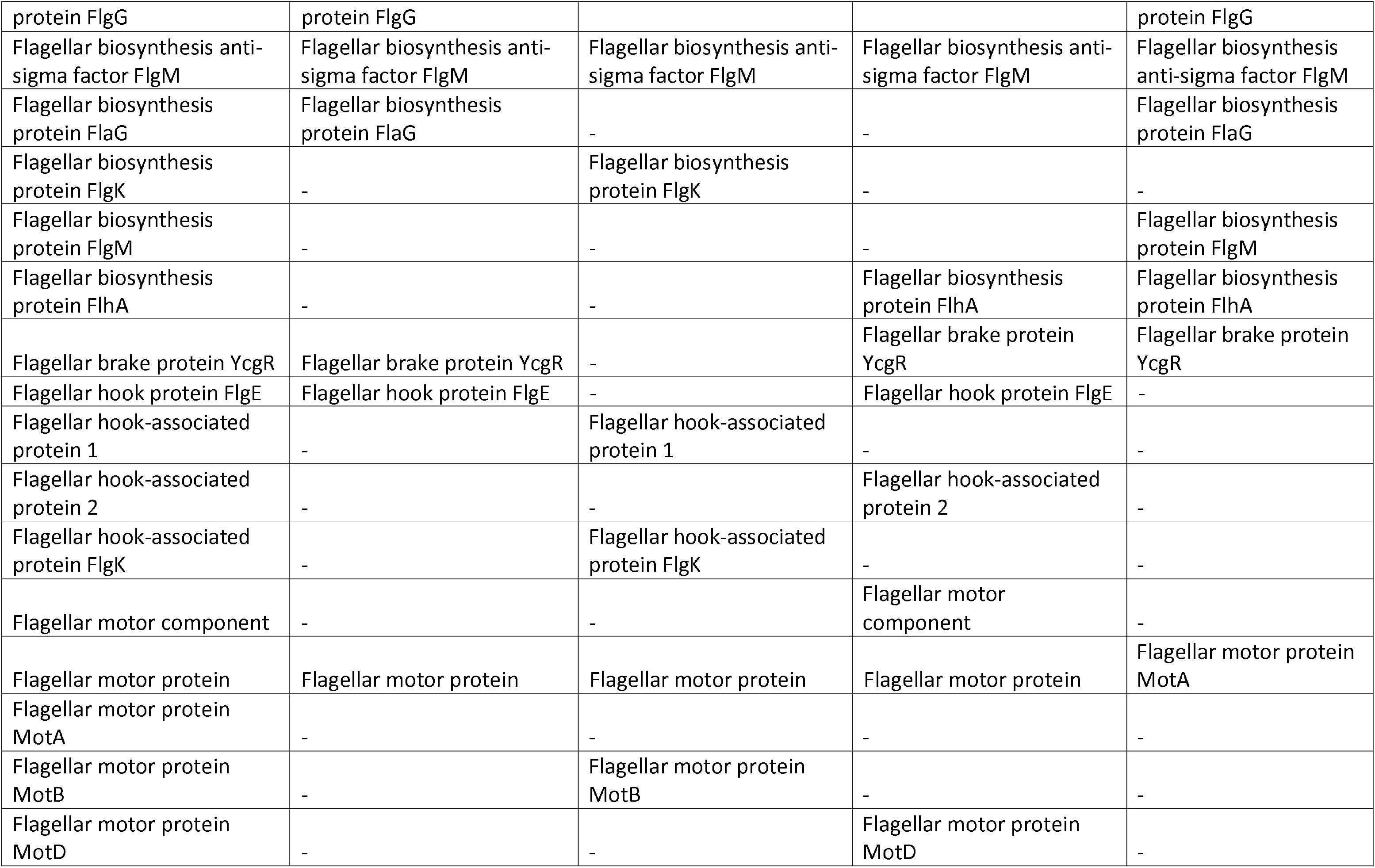
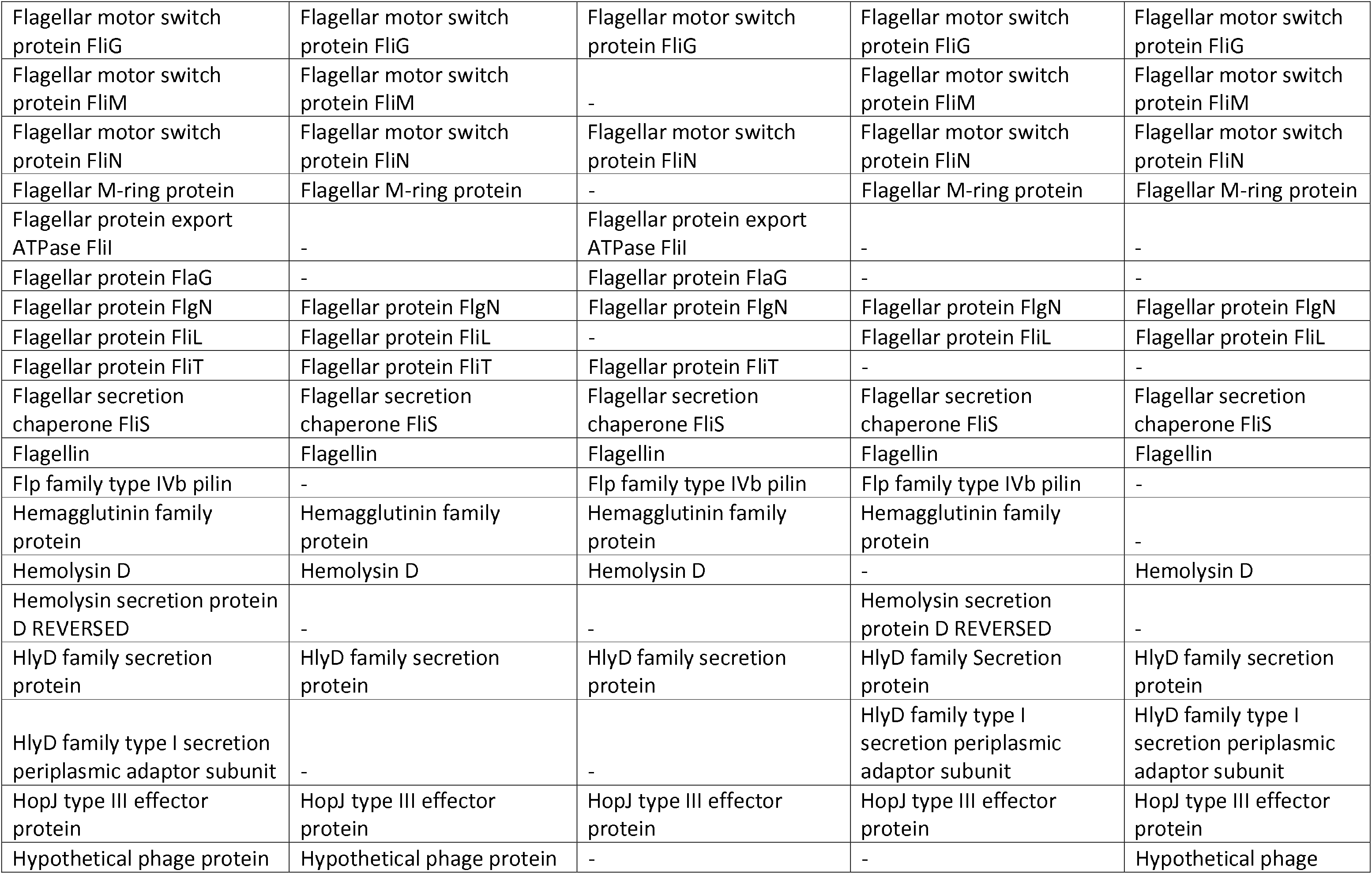
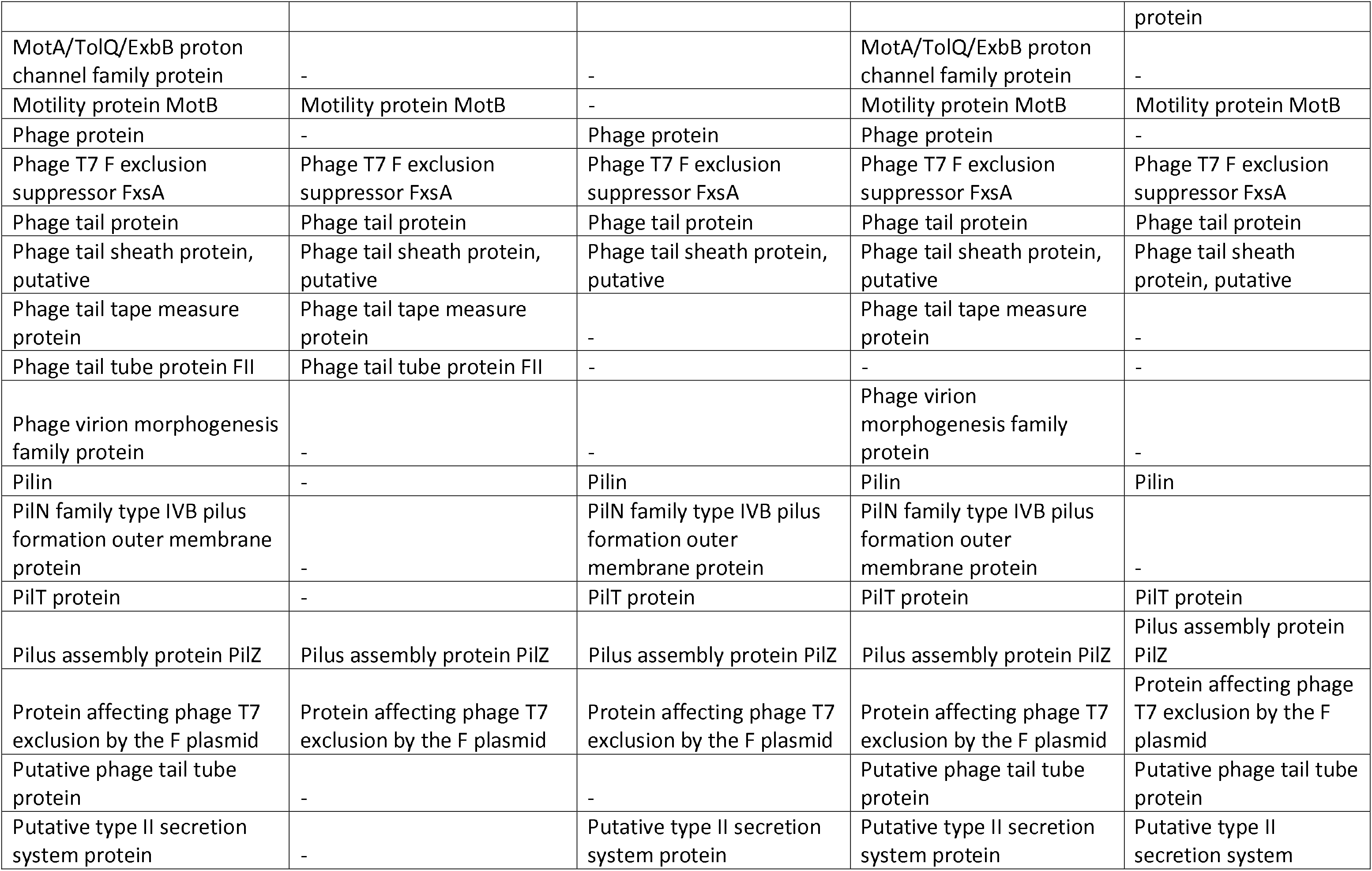
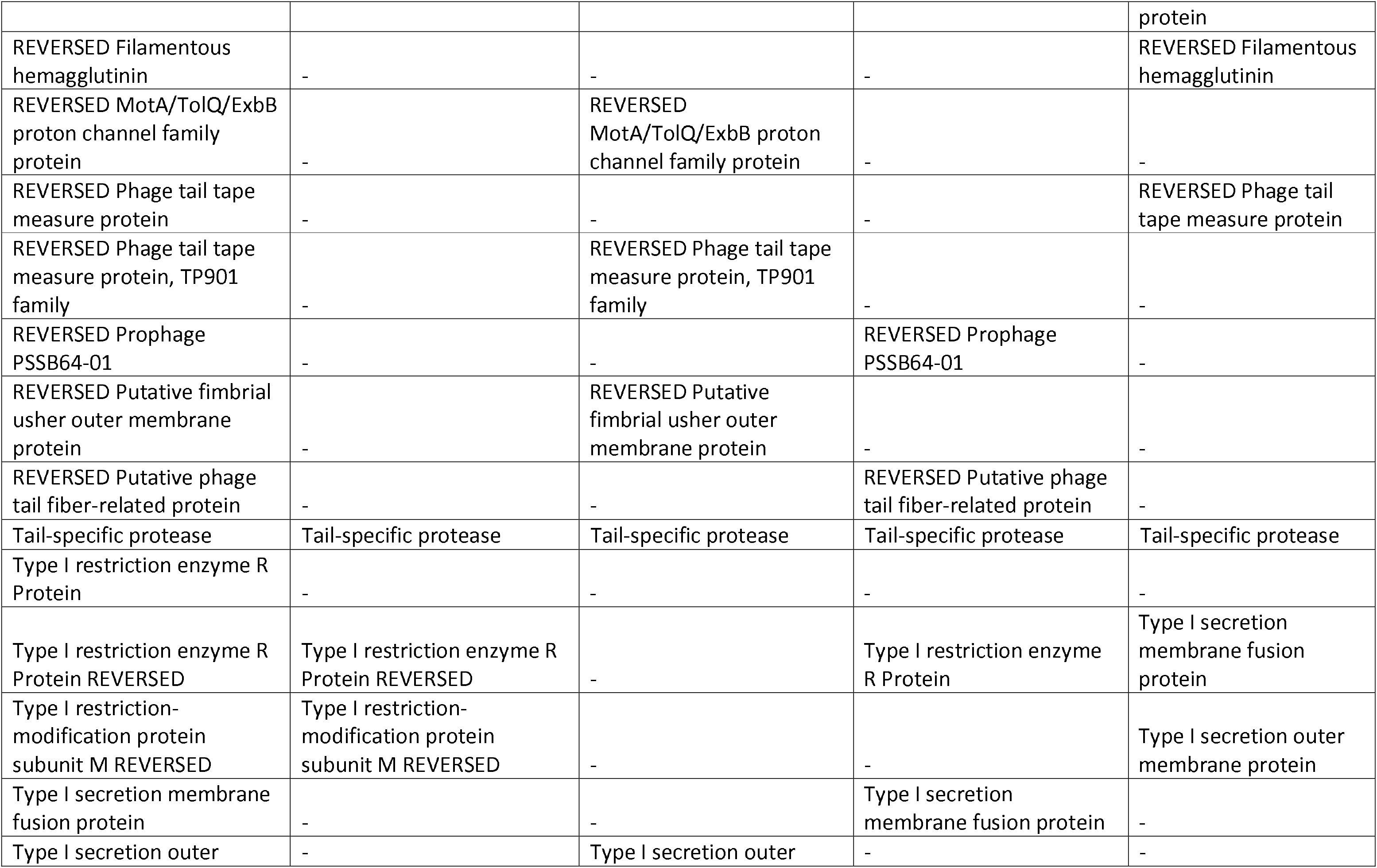
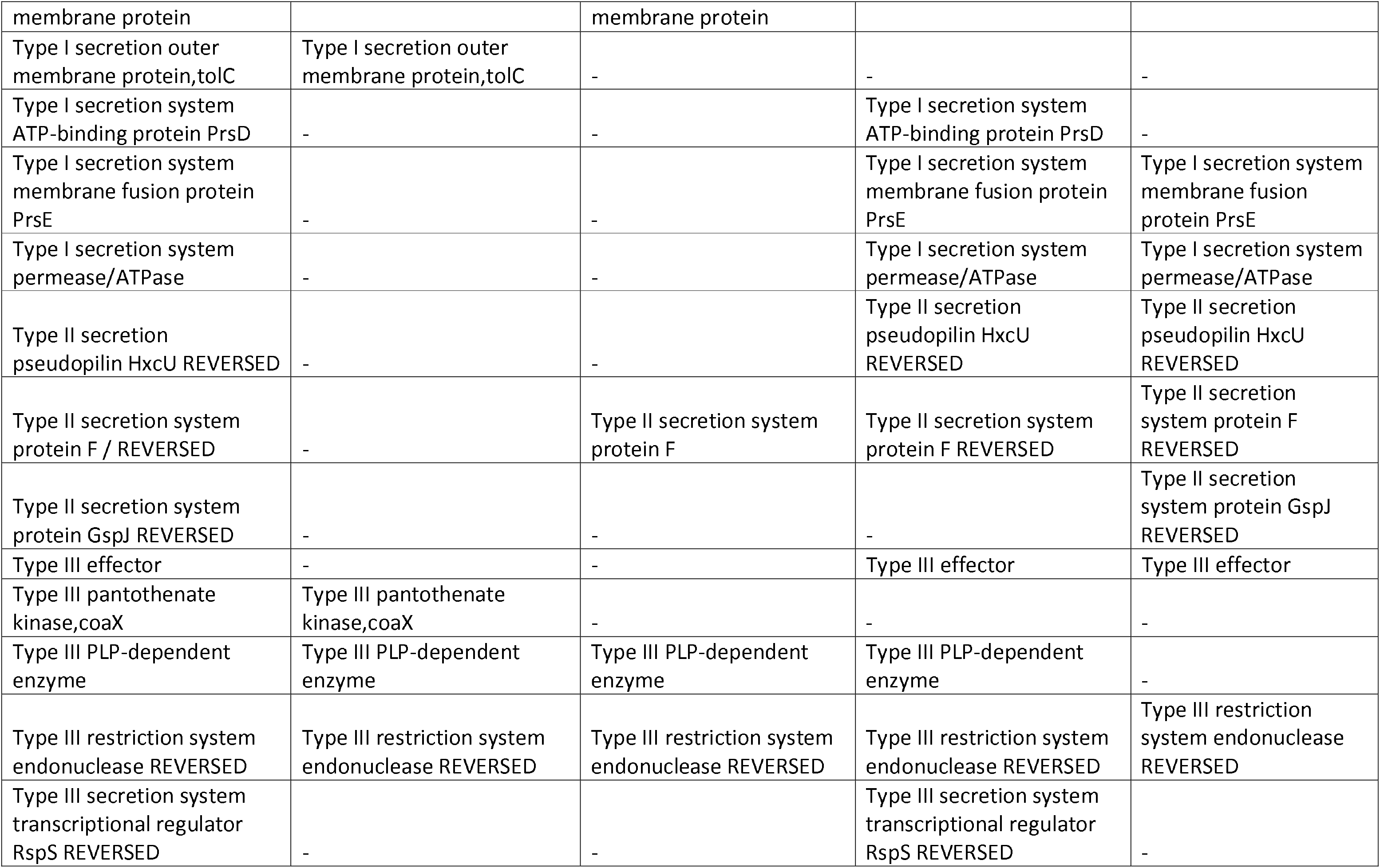
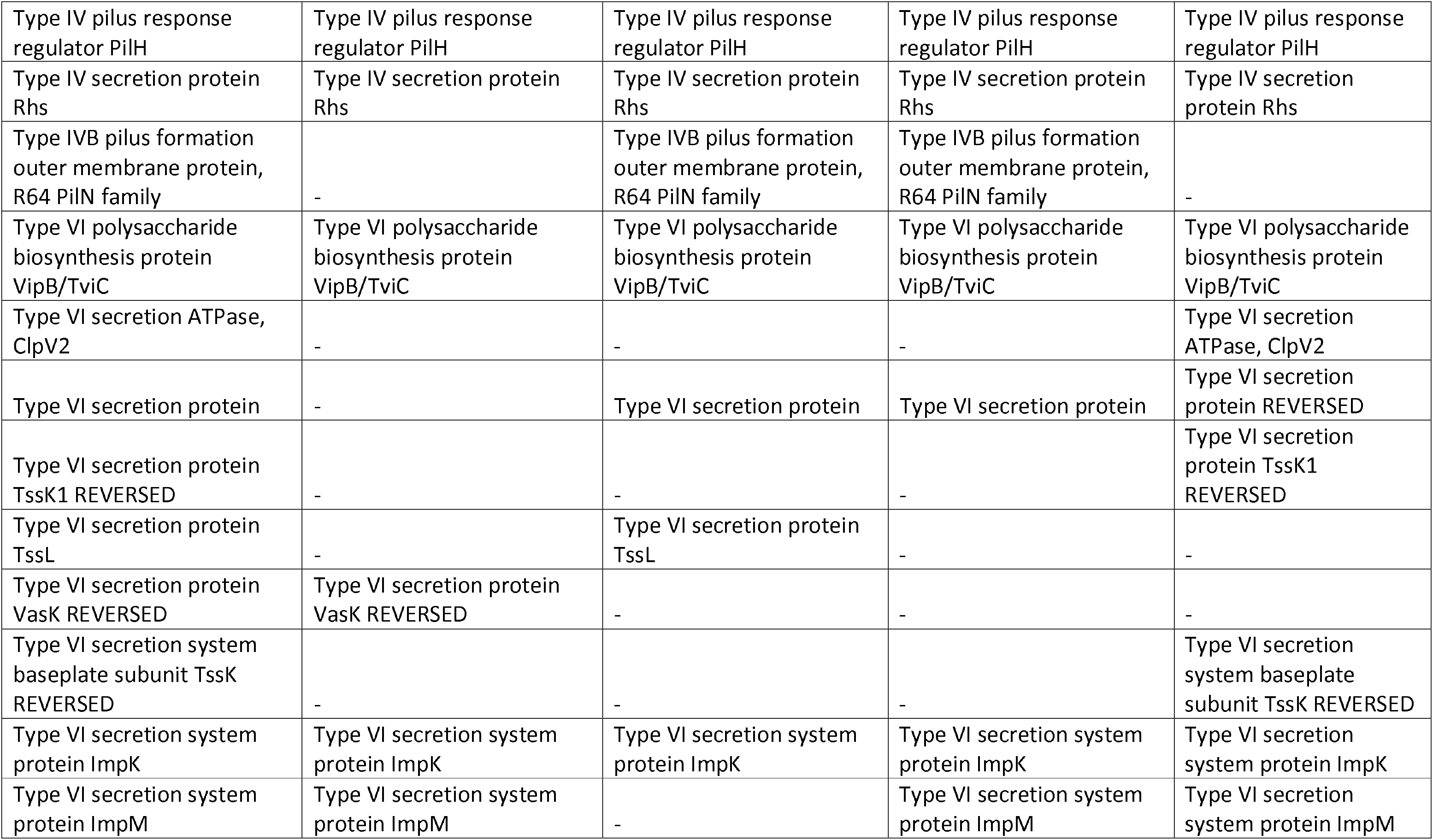
Protein summary of the four *Pseudomonas fluorescens* isolates: This table provides the details of different secretion systems, fimbria and flagella related proteins, phage and phage related proteins and hemolysin proteins present in all four isolates.

**Supplementary Table 2:** Protein modifications and peptide summaries of PFR46H06, PFR46H07, PFR46H08 and PFR46H09. This table summarizes proteins that have undergone amino acid modifications, the positions where protein modifications have taken place and other chemical modifications based on the mass-spectra of those proteins. See the attached Excel sheet.

## Supplementary Figures

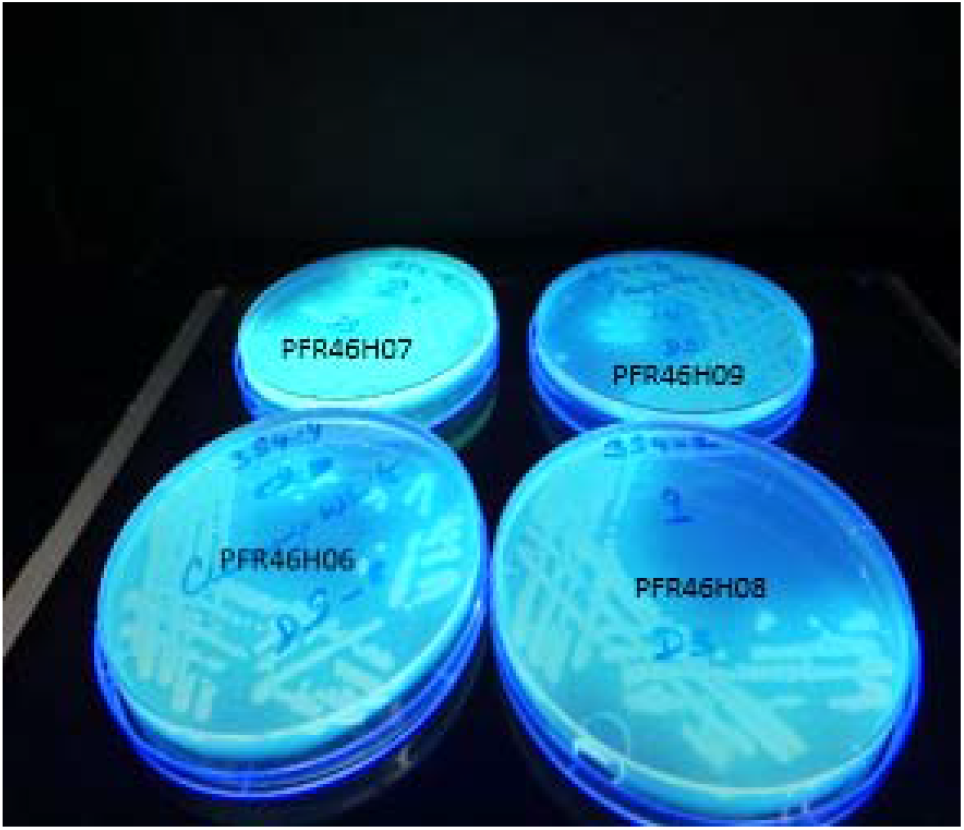
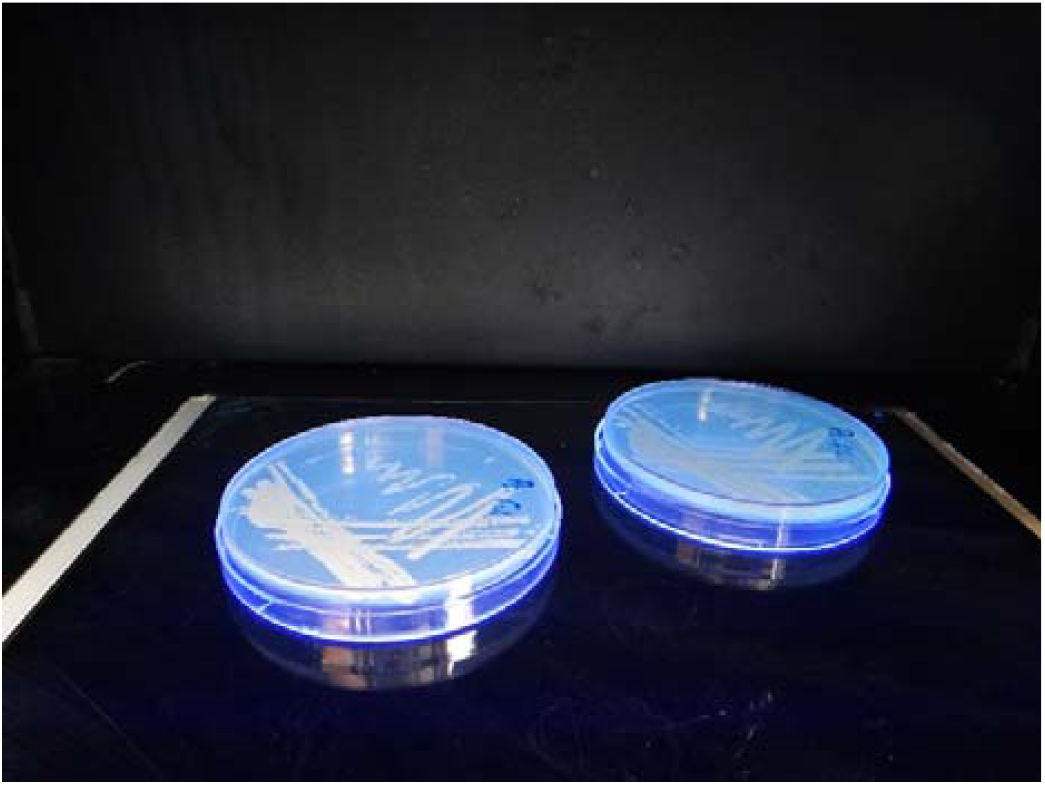
Supplementary Figure 1a shows the four strains of *Pseudomonas fluorescens* cultures on BHI agar plates PFR46H06, PFR46H07, PFR46H08, PFR46H09 fluorescing under UV illumination. Figure 1b shows the duplicate plates of PFR46I06 where the colonies did not show conspicuous florescence.

**Supplementary Figure 2:**
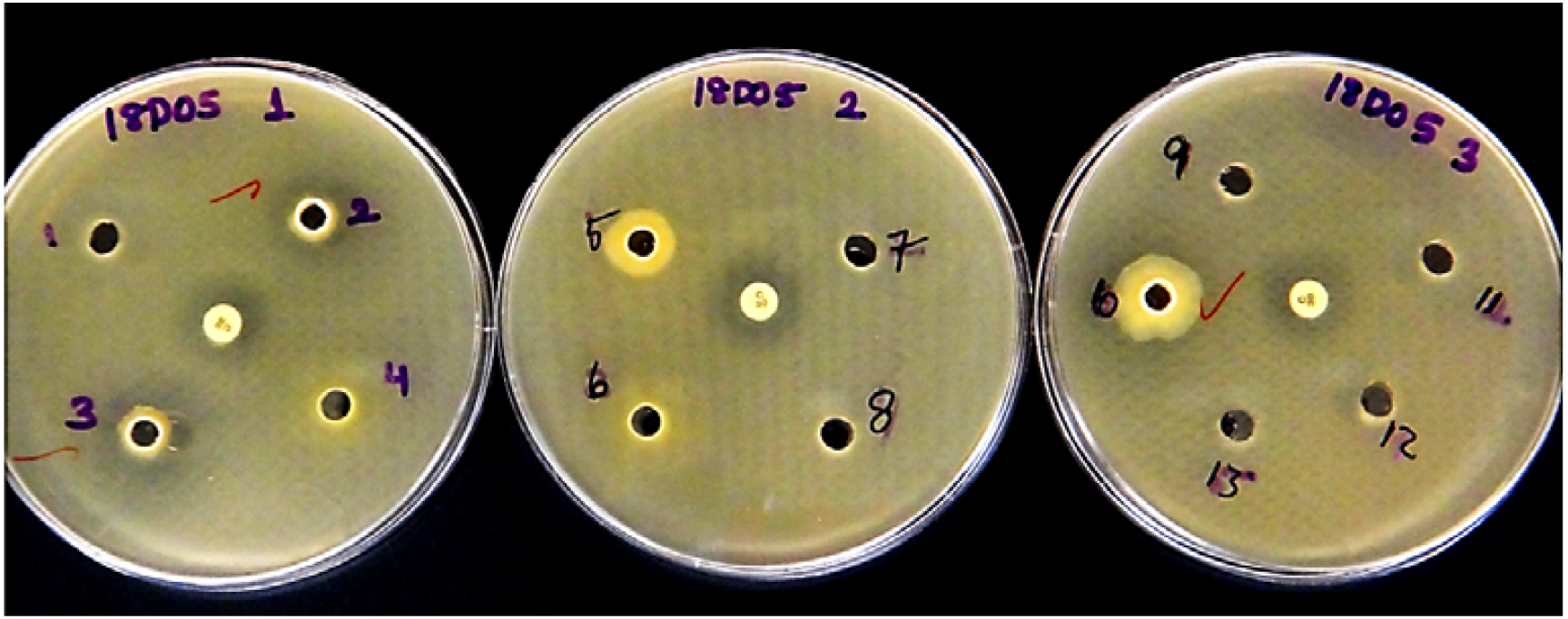
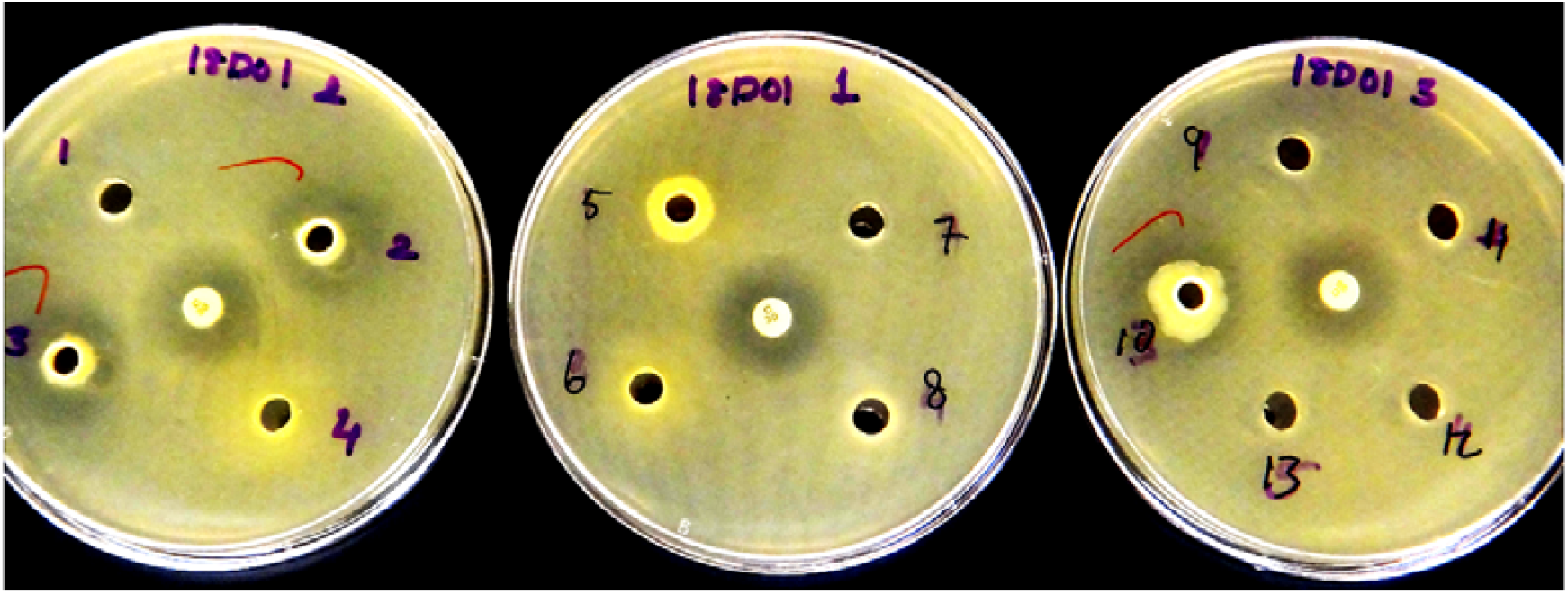
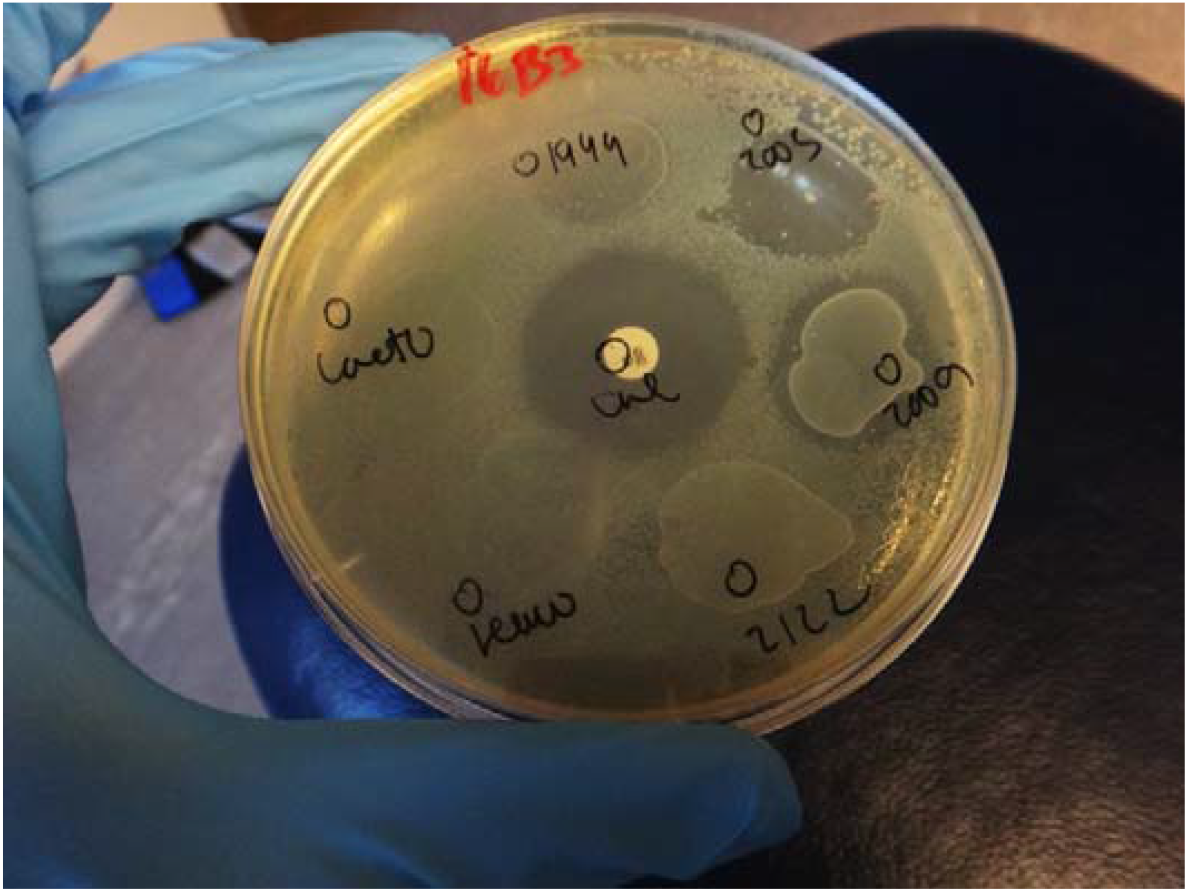
Zones of inhibition produced by five *Pseudomonas fluorescens* strains against *Listeria monocytogenes* in pour plates. 2a: After 48 h at 30°C. 2b: After 48 h at 20°C. 2c: Drops of unfiltered supernatants from proven protective cultures. Wells numbered 2, 3, 4, 5 and 10 (Figures 2a and 2b) correspond to cultures PFR46H07, PFR46H08, PFR46H06, PFR46I06 and PFR46H09 respectively. Zones labelled 1944, 2003, 2009, 2122, Leuco and Lacto (Figure 2c) correspond to supernatants from *Carnobacterium maltaromaticum* 1944, 2003, 2009, *Carnobacterium divergens* 2122, *Leuconostoc gelidum,* and *Lactococcus piscium* respectively. Chloramphenicol discs (30 μg) are also in the center of each plate as a positive controls.

## Notes

http://www.ebi.ac.uk/pride

https://drive.google.com/drive/folders/1ZlbR6UaR0kC5LnZymSXj3ZuSl8qdavyN

### Competing Interest Statement

The authors have declared no competing interest.

